# Assessing structure-function impacts on Vitellogenin by leveraging allelic variant occurring in honey bee subspecies *Apis mellifera meliffera*

**DOI:** 10.1101/2025.03.17.643649

**Authors:** Vilde Leipart, Oriol Gracia I Carmona, Christine Orengo, Franca Fraternali, Gro V Amdam

**Affiliations:** Faculty of Environmental Sciences and Natural Resource Management, Norwegian University of Life Sciences, Ås, Norway; Department of Structural and Molecular Biology, University College London, London, UK; Randall Centre for Cell & Molecular Biophysics, King’s College London, New Hunt’s House, Guy’s Campus, London, UK; School of Life Sciences, Arizona State University, Tempe, Arizona, USA

**Keywords:** Honey bee vitellogenin, genetic diversity, structural bioinformatics, molecular dynamics simulations, protein language model variant prediction, structure-function relationships

## Abstract

Computational advances involving artificial intelligence (AI) and successful experimental state-of-the-art structure determination can provide detailed pictures of large and complex protein structures, and their variations. A standout case is Vitellogenin (Vg) derived from the honey bee (*Apis mellifera*). Vg is an essential protein for reproduction in almost all egg-laying animals, and can in addition regulate behavior and provide immunological support in some species, including the honey bee. Information is limited in terms of the structure-function relationships that underlie Vg’s pleiotropic functions, at least in part because this protein is not expressed in the best developed gene-editing models, such as fruit flies and mice. However, naturally occurring allelic variation in Vg can provide some insight, i.e., into which changes (mutations) are allowable (present at some frequency) vs. likely not allowable (not present or only at low frequency). Here, we leverage a unique dataset of 1,086 fully sequenced Vg alleles from honey bees in 15 countries. We identify a population-specific 9 nucleotide deletion in a locally endangered honey bee subspecies (*A. m. mellifera*) that impacts a loop structure in a central Vg domain. Due to the *A. m. mellifera* population history of near extinction and human intervention, an assessment of this Vg variant is not only theoretically interesting but also relevant for subspecies conservation efforts. Using structural bioinformatics, molecular dynamics simulations, and a transformer-based indel predictor (IndeLLM), we demonstrate that Vg protein structure and stability can be maintained despite the deletion. Our approach also reveals the dynamic nature of specific regions in Vg for the first time. Generalizable results may extend to other egg-laying animals of ecological and economic importance.

## Introduction

Solving the structure of complex biological molecules has challenged researchers for decades. Large sizes, flexible regions, and hydrophobic surfaces have hindered traditional experimental structure determination. The recent revolution in artificial intelligence (AI) has begun a new area where structural puzzles can be predicted with remarkable accuracy [1,2]. Also, deep learning methods help to understand the function and dynamics of proteins by predicting the relationship between the sequence and structure [3]. An example of a challenging biological molecule is the egg yolk precursor protein Vitellogenin (Vg), which is broadly present in egg-laying animal taxa. Vg is a large, multifunctional protein, best known for being essential for female reproduction [4], but it can also regulate complex behavior in some insects [5–7], and provide immunological functions in coral [8], scallop [9], crab [10], insects [11,12] and fish [13,14]. The first structure determination of Vg was resolved close to 25 years ago from a Lamprey (*Ichthyomyzon unicuspis*)[15]. This crystal structure mainly covered functional regions (75% of the protein sequence) responsible for binding and transporting nutrients to developing embryos. The remaining protein sequence could not be resolved. Discovering the full-length structure of Vg and expanding the structural understanding to more species has been a slow process.

The recent computational advances involving AI and deep learning have provided access to high quality predicted protein structures. The AI-based AlphaFold 2 (AF) algorithm was the first method to achieve a prediction accuracy comparable to experimental methods [1]. When AF was released, a database of its predictions was also made publicly available (AFDB [16]). AFDB quickly became a central data resource with over 200 million structures [16], including close to 2,000 Vg structures spanning diverse species and groups, including chicken, fish, geckos, nematodes, and insects. In this context, contributions made by structural work on honey bee (*Apis mellifera*) Vg has been highlighted as a prime example where AF is able to produce high quality models (confidence score, pLDDT, > 80) [17– 19]. Vg represents an exposé of contextual and molecular properties that showcase the impacts of AI-driven structural determination: The biologically and economically important protein is of a large molecular size is responsible for complex, pleiotropic traits involved in health and behavior. It has multiple interacting domains, complex post-translational modifications, metal-, ligand- and or receptor-binding capabilities, considerable capacity for cargo, alternative cleavage products, and the list goes on [4,6,14,20–34].

On top of computational advances, large strides have also been taken experimentally to understand honey bee Vg structure. The native protein was recently resolved using cryo-electron microscopy (cryo-EM), while the AF2 model supported model refinmet, the experimental density confirmed an impressively accurate prediction by AF (RMSD: 2.35 Å [24]). Also, the high-resolution cryo-EM map shows non-protein molecules and cleavage products. The AF model does not capture such features. Therefore, we have obtained a detailed structural understanding of honey bee Vg by combining the information from both models: a large multidomain structure, including 3 conserved domains typically found in protein family members, but also includes 2 domains of unknown molecular function [23,24]. The structural representations further identify multiple disordered and potentially flexible regions, a large hydrophobic cavity, several possible proteolytic cleavage sites, a post-translational modification, and metal binding sites [23,24,28,35].

Taken together, current molecular models provide good insight into several layers of complexity for Vg. Yet, these models account for only one variant of the protein: The AF model was built from the honey bee Vg reference sequence (UniProt ID: Q868N5), and the cryo-EM model relied on the AF model for map quality and model building. This situation does not adequately reflect the genetic variantion in honey bees as a species [36–38]. In fact, we previously discovered 121 different versions (variants) of the Vg protein in an allelic survey of honey bees from 15 countries [38]. Such alterations in amino acid composition can impact the protein’s fold, stability, dynamics, and, consequently, its functions.

To gain a more accurate structural understanding of Vg, it is valuable to concider the sequence variation that occurs in the protein. What type of substitutions, deletions, or additions are present, their distribution in the protein structure and frequency.

The impact of amino acid substitutions, deletions, and additions can be studied experimentally in protein model systems, most recently by leveraging CRISPR-CAS9 technology. However, large and complex proteins like Vg are challenging to target comprehensively in such analyses, and also, many species that rely on this protein are not well-developed models. In this context, the availability of 1,086 full-length allelic sequences for honey bee Vg is valuable recouce [38]. The identified variation is a snapshot of naturally established variation resulting from random mutations and various selective forces. Any variant present at a reasonable frequency, moreover, is unlikely to confer disastrous organismal consequences under normal conditions. With a long list of natural variation as a starting point, we can start probing Vg variants to understand the flexibility of Vg regarding what the protein can allow, and not allow, in terms of changes of type and number of amino acids.

The first studies looking into the naturally occurring variation in honey bee Vg found that the C-terminal half of the protein-coding sequence was the main site for diversity [36]. This region encodes the nutrient transport cavity, primarily responsible for carrying lipid molecules and metal ions. More recently, we compiled a new dataset from a broader geographical range and with the use of sequencing technology that allowed entire alleles to be assembeled [38]. When analyzing those data, we identified that non-synonymous (coding) single nucleotide polymorphisms (nsSNPs) were distributed non-uniformly across the domains of Vg [38]. We confirmed that the lipid-binding cavity is highly enriched in mutations, while the N-terminal β-barrel is highly conserved [23,24,33,38]. Unlike the lipid-binding cavity, the β-barrel appears to have multiple functional roles, containing a receptor recognition site [39–41], a proteolytic cleavage site [25], one or more zinc (Zn) binding sites [28], the capacity to bind DNA [29] as well as a providing a site for post-translational modifications such as glycosylation and phosphorylation [24,25,31].

Yet, despite the conserved nature of the β-barrel, our dataset of allelic variation pointed to the presence of a short deletion in this region of Vg. This deletion is highly frequent in one population: the European Dark Bee subspecies (*A. m. mellifera*), which is classified as locally endangered [42,43]. During the 1990s, several initiatives emerged to protect the gene pool of the European Dark Bee [44], leading to strategies in which native Dark Bees were sheltered in their environment by creating conservatory apiaries [45–50]. The observation of a deletion in Vg that is mainly confined to honey bees with this conservational history might raise concerns about the effect of the mutation, and whether or not it is present because of low effective population sizes or other unintended and detrimental effects of closed breeding. Alternatively, the deletion could be a neutral or even beneficial change – a unique population-specific isoform.

In this article, we leverage our dataset of Vg allelic variation to study the structural impacts of the deletion observed in the Vg β-barrel of *A. m. mellifera*.

In studying these impacts, we had to factor in that the dynamical features of the β-barrel are not understood, and there is generally little detailed knowledge about the atomic mechanisms during Vg’s proteolytic cleavage or interactions with DNA, receptors, and zinc. Thus, we chose to utlize the toolbox of molecular dynamics and an inhouse developed indel pathogenicity predictor [51] to predict if the deletion confers any structural or fuctional impacts on the β-barrel.

This is the first study of its kind to reveal the dynamic nature of structural regions in Vg. Our findings suggest that the protein structure and stability are maintained despite the deletion. We further reveal that the Vg β-barrel has unique structural features despite the high conservation across the protein family, and that these features have neutral structural consequences. Our study contributes to understanding a locally endangered subspecies’ genetic pool and value.

## Results

### Deletions in the coding region of the *vg* gene

We have identified three different-sized deletions in the exon regions of the *vg* gene: p.N153_V155del, p.S844_V845del, and p.R1669del (Fig. 1A). The deletions are located in exons 2, 4, and 7, respectively, and were identified at varying frequencies (Fig 1B-1D). The p.N153_V155del deletion is located in the β-barrel and was found in 105 haplotype sequences in total, with a majority (91 sequences) coming from *A. m. mellifera* conservatory apiaries in Europe (Fig. 1B). Of the 9 conservatories we surveyed, the p.N153_V155del deletion was identified in 8 of them, and our samples included 16 homozygotic individuals (Fig S1). In contrast, the two deletions outside the β-barrel (exons 4 and 7) were found at single apiaries in the USA in 3 and 1 haplotype sequences, respectively (Fig 1C-1D), and no homozygotic individuals were identified. Taken together, this information suggests that only the p.N153_V155del deletion is present at a frequency that clearly makes it biologically relevant and interesting.

**Figure 1.**
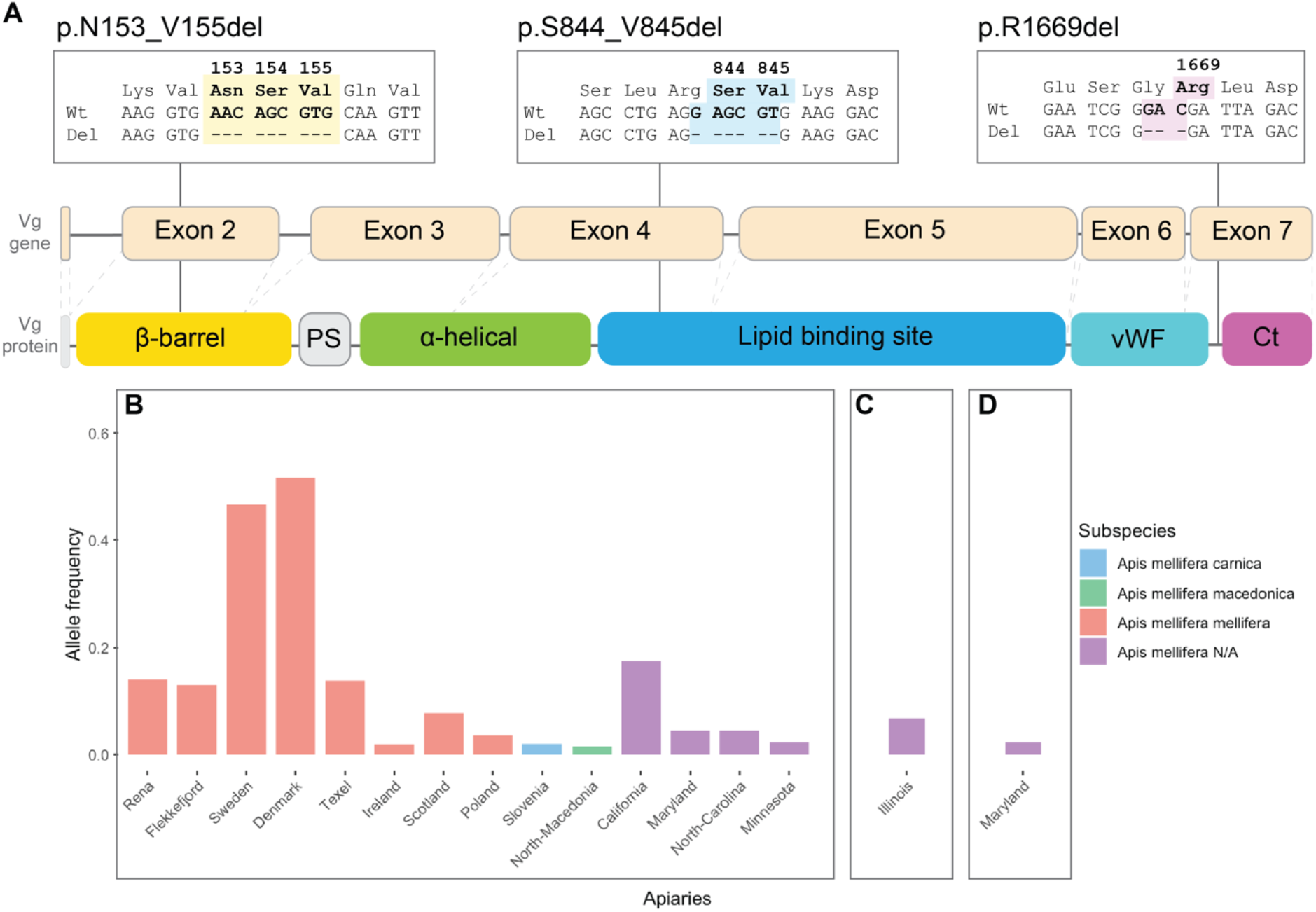
The identified deletions and their allele frequencies: **A** The three boxes shows a small sequence alignment for the identified deletions: the amino acid (aa) numbers, aa types, the wildtype (wt) nucleotide sequence, and the deleted (del) nucleotide sequence. The deleted region is shown in bold text and colored yellow (p.N153_V155del), blue (p.S844_V845del), or pink (p.R1669). The boxes are positioned above a 2D representation of the vg gene (exons numbered) and a 2D representation of the Vg protein, and the line from the alignment box show where the deletions occur in the gene and protein. The Vg protein include the signal peptide (white), the β-barrel (yellow), the polyserine linker (PS, grey), the α-helical (green), the lipid binding site (blue), the von Willebrand factor domain (vWF, cyan) and the C-terminal region (Ct, magenta). **B-D** The allele frequency (y-axis) per apiary (x-axis) for the three deletions: p.N153_V155del in panel B, p.S844_V845del in panel C, and p.R1669 in panel D. The apiaries are colored by subspecies (legend). In panel B, the haplotype sequences from Slovenia, Notrh-Macedonia, Minnesota, and one from Maryland are idenfied as recombinant haplotypes, which leaves only 10 haplotype sequences identified outside the conservatories for A. m. mellifera.

Translating the Vg gene sequence including the p.N153_V155del deletion shortens a loop in the β-barrel by 3 amino acids. In theory, reducing the length of a loop could improve the stability of a protein fold [52,53]. However, a shorter loop could hinder movement, which could be problematic if the protein requires flexibility to perform its task. We thus decided to explore whether the loop shortening may impact β-barrel structure, stability, or dynamics.

### The long deletion loop in the β-barrel is not conserved and exposed

To evaluate whether the deletion is in a conserved region, we built a functional family to identify the residues with functional importance (see methods for details). The final multiple sequence alignment (MSA) was used to calculate a conservation score for each residue in the β-barrel (Fig. 2A). The deleted residues (aa 153-155) are located in an unconserved loop (aa 141-164, average conservation score of 0.02 out of a possible max of 1.00). Also, the deletion loop is on the surface of the β-barrel facing in the opposite direction to the lipid binding cavity (2B). The deletion loop is 19 residues long and contains a short α-helix in the middle of the loop. The deletion occurs within the α-helix (Fig. 2C).

**Figure 2.**
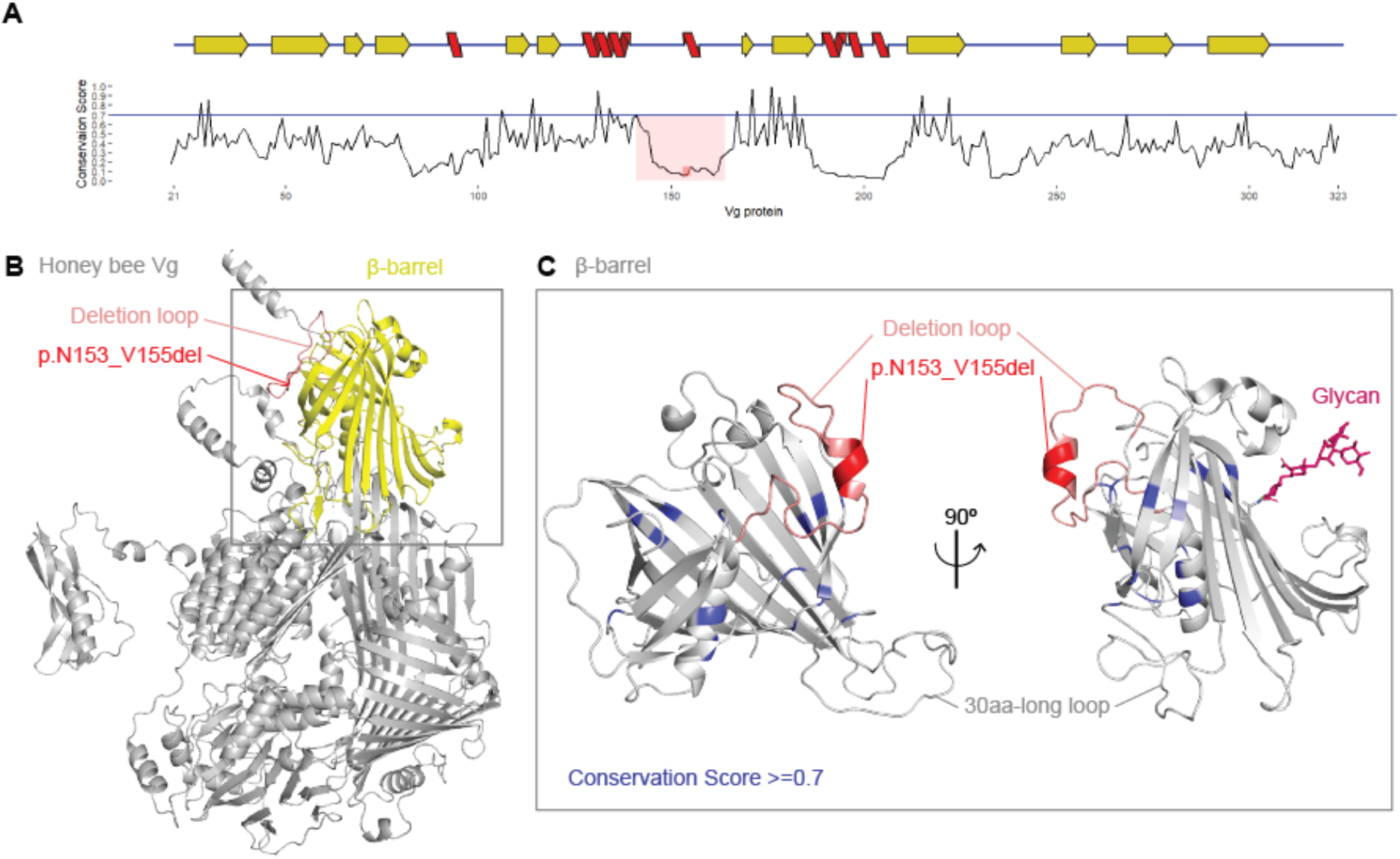
Conservation and structural position: **A** The conservation score (y-axis) is plotted for the aa in the β-barrel (x-axis, aa 21-323). The threshold for functionally conserved is shown as a blue line (0.7). The deleted residues are in the red box (aa 153-155), while the deletion loop is in the pink box (aa 141-167). Above the plot is a 2D representation of the secondary structure elements in the β-barrel (β-strands are yellow, while α-helices are red). **B** The β-barrel (yellow) is the first structural region in the full-length honey bee Vg structure (grey). The deletion loop and deletion are labeled as in panel A. **C** Two orientations of the β-barrel (grey) are shown, and conserved sites are colored blue (>0.7). The deletion loop and deletion are colored as in panel A. The glycan is shown on the orientation to the right (pink), and the 30aa-long loop is labeled.

### Generated four models of the β-barrel to account for glycosylation impact and flexibility

To investigate whether the deletion could have an effect on the structure stability, we performed molecular dynamics simulation of a wildtype and deleted system. A Cryo-EM model of honey bee Vg identified N-linked glycosylation on N296 with unknown structural or functional importance (Fig 2C)[24]. In general, N-linked glycosylation impacts proteins’ physical properties and biological functions [54]. Therefore, it was important to include the glycan in our simulations. However, glycans can be highly dynamic. To separate any effect contributed by glycan flexibility, we generated four systems for molecular simulations: 1) wildtype (wt), 2) wildtype with glycan (wtg), 3) deletion (del), and 4) deletion with glycan (delg). We performed 60 μs of molecular dynamic simulation (15 μs of aggregated data per system) using five replicates of 3 μs per system. The first 400ns were removed for the following calculations based on the root mean square deviation (RMSD) profile convergence (Fig S2).

### The deletion does not affect the stability to the β-barrel

The radius of gyration (Rg) was calculated for each replicate in the four systems. The Rg for the wt starting structure was 20.12 Å, and the Rg measured for all the replicates over the entire simulation stayed within the range from 19.06 to 21.60 (Fig S3A-S3D). We concluded that the protein backbone compactness remained constant through the simulations for all the replicates, and there were no differences between the systems (Fig 3A). Next, we calculated the mean RMSD of the Cα atoms for the four systems and found no significant difference (Fig.3B and Table 1 for mean RMSD and see Fig S3E-S3H for variation between replicates). Overall, we concluded that neither the deletion nor the glycosylation resulted in significant structural differences or instabilities of the overall structure of the β-barrel.

**Table 1:**
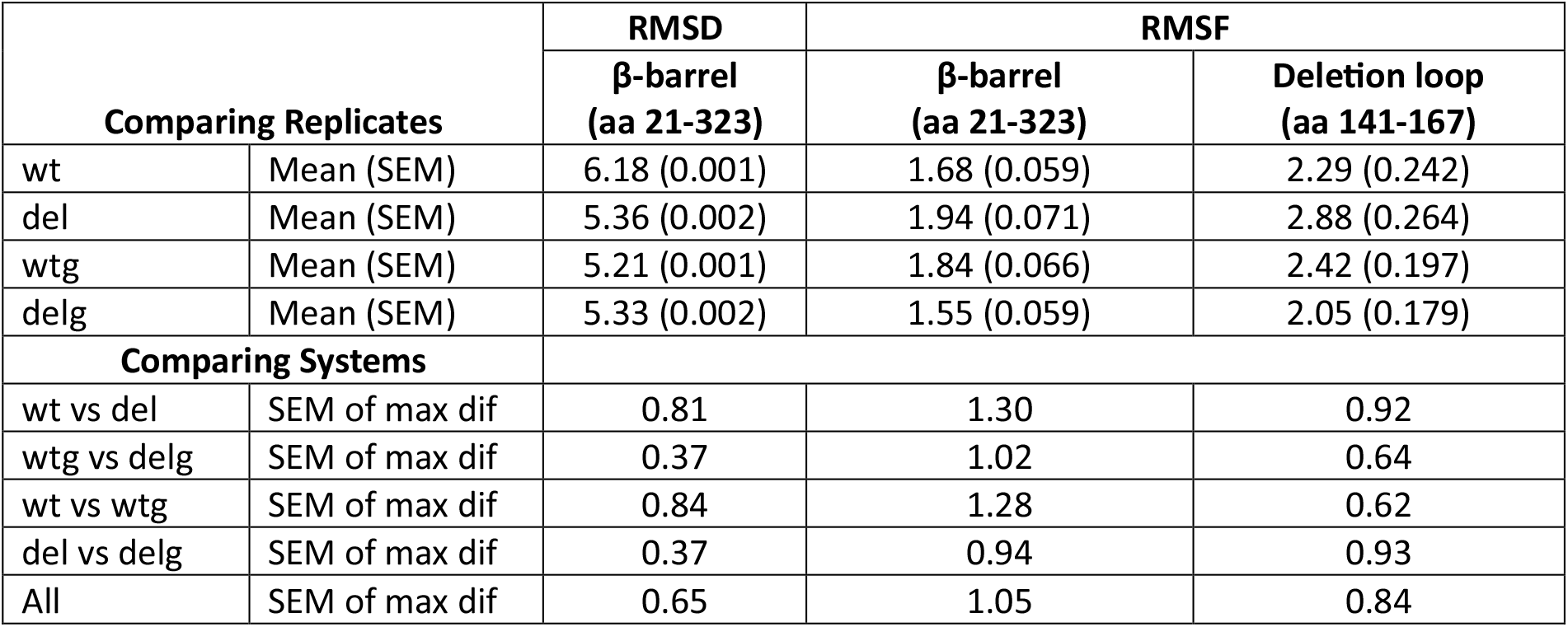
RMSD and RMSF calculations. Rows 1-4 show the calculations for RMSD and RMSF within the replicates for the systems (wt, del, wtg and delg). We calculate the overall mean and standard error of the mean (SEM) for the β-barrel, but also for the full and short deletion loop. The Max dif refers to the time frame (ns for RMSD) or position (Cα for RMSF) where we observed the maximum difference between the systems using the mean values from the systems, where we calculate the SEM between the systems.

| Comparing Replicates |  | RMSD | RMSF |  |
| --- | --- | --- | --- | --- |
| | | β-barrel<br>(aa 21-323) | β-barrel<br>(aa 21-323) | Deletion loop<br>(aa 141-167) |
| wt | Mean (SEM) | 6.18 (0.001) | 1.68 (0.059) | 2.29 (0.242) |
| del | Mean (SEM) | 5.36 (0.002) | 1.94 (0.071) | 2.88 (0.264) |
| wtg | Mean (SEM) | 5.21 (0.001) | 1.84 (0.066) | 2.42 (0.197) |
| delg | Mean (SEM) | 5.33 (0.002) | 1.55 (0.059) | 2.05 (0.179) |
| Comparing Systems |  |  |  |  |
| wt vs del | SEM of max dif | 0.81 | 1.30 | 0.92 |
| wtg vs delg | SEM of max dif | 0.37 | 1.02 | 0.64 |
| wt vs wtg | SEM of max dif | 0.84 | 1.28 | 0.62 |
| del vs delg | SEM of max dif | 0.37 | 0.94 | 0.93 |
| All | SEM of max dif | 0.65 | 1.05 | 0.84 |

**Figure 3.**
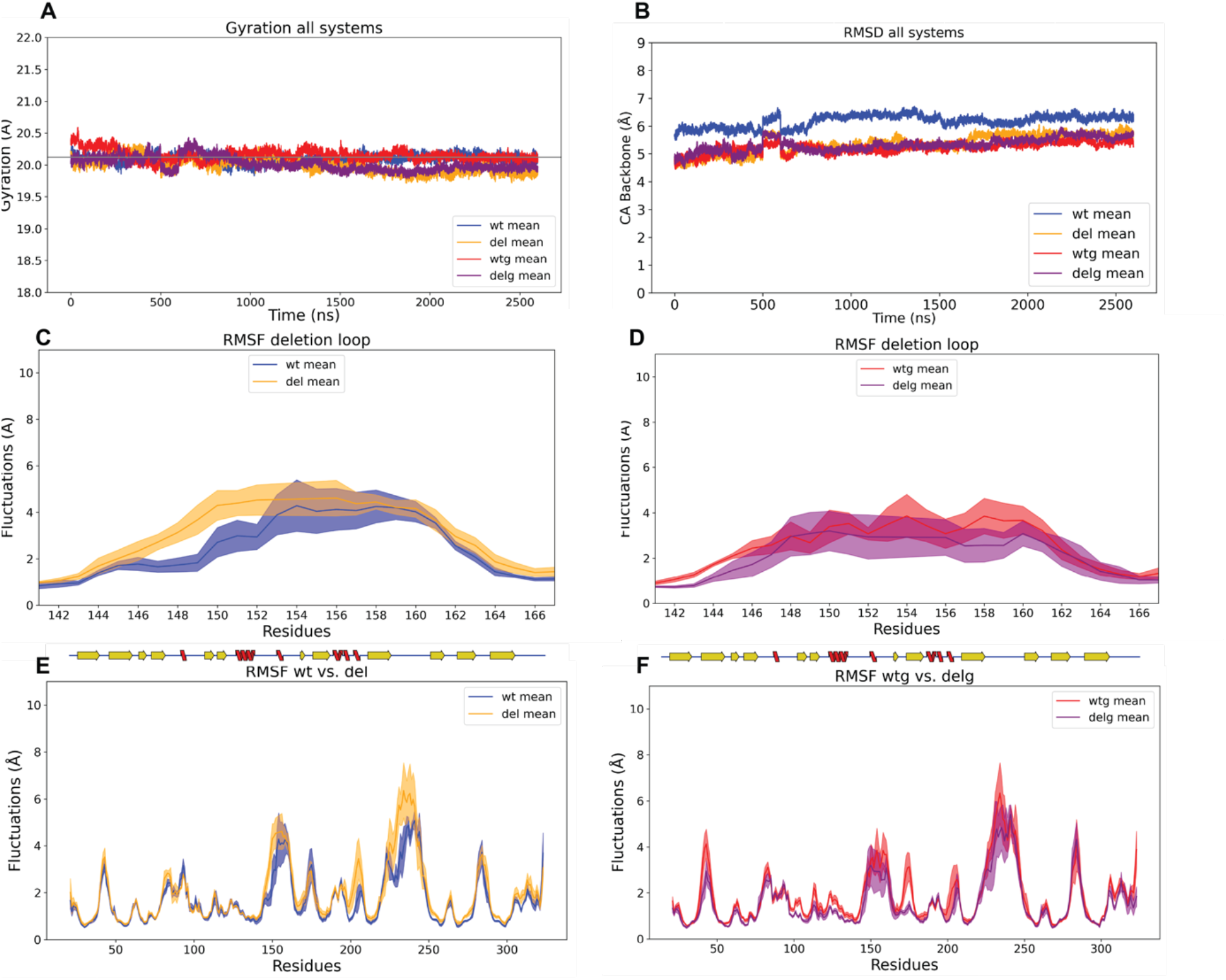
Rg, RMSD, and RMSF: **A** The mean Rg (y-axis) over time (x-axis) for each system (wt: blue, del: orange, wtg: red, and delg: purple). The Rg for the wt starting structure (20.12 Å) is included as a grey line. **B** The mean RMSD between the Cα atoms (y-axis) over time (x-axis) for each system. Same colors as in panel A. **C** The mean RMSF (y-axis) for the deletion loop Cα in the wt (blue) and del (orange) systems (x-axis, aa 141-167). The mean values are plotted as a solid colored line, while the ± SD values are the same color, but transparent. **D** Same plot as panel C, but the mean RMSF values are from the wtg (red) and delg (purple) systems. **E** The mean RMSF (y-axis) for each Cα in the wt (blue) and del (orange) systems (x-axis, aa 21-323). The mean values are plotted as a solid line, while the ± SD values are transparent. On top of the plot is the 2D representation of the β-barrel secondary structure elements, the same as in Fig 1A. **F** has the same type of plot as panel C, but the mean RMSF values are from the wtg (red) and delg (purple) systems.

### No difference in the flexibility of the deletion loop

To highlight differences in the flexibility of individual backbone atoms, we calculated the root mean square fluctuations (RMSF) of the Cα atoms per replicate and the mean RMSF per system (Fig S4A-S4D). The most fluctuating Cα atoms in the del, wtg, and delg systems were within a 30-amino acid (aa) loop region (grey loop in Fig 2C), while a Cα in the deletion loop fluctuated the most in the wt system (pink loop in Fig 2C). This suggests that the flexibility of the deletion loop is higher for the wildtype than for the deleted or glycosylated systems, but these differences are not significant (T-test, wt vs. del: p = 0.275, wt vs. wtg: p = 1.000 and wtg vs. delg: p = 0.380). We expected that shortening a loop could reduce the range of motion for the deletion loop and, therefore, the deleted systems could have significantly lower flexibility compared to the wildtype systems, but comparing the mean RMSF observed for the deletion loop (Fig. 3C-3D and Table 1) between the wt and del in both non-glycosylated and glycosylated systems, we found no significant differences (T-test, wt vs. del: p = 0.107 and wtg vs. delg: p = 0.174). In addition, the resolution of our starting model is 2.5-3.0 Å [24], and therefore, changes smaller than our resolution is within the margin of error. Taken together, we concluded that the minor differences in the flexibility of the deletion loop result from replicate variability, and we observe no consistent difference between the wt and del systems.

### No conformational effects caused by the deletion were observed

We compared the mean wt RMSF to the mean del RMSF of the complete backbone to identify if the deletion affected the protein conformational stability. Plotting the mean values (with ± 1 SD) showed an almost complete overlap (Fig. 3E-3F). To identify potential significant differences caused by the deletion (wt vs. del and wtg vs. del), we compared the mean RMSF of Cα in groups of 3 (Cα peaks) across the entire backbone (sliding window with an increase of 1). The significant Cα peaks are listed in Table 2, calculated using a t-test (alpha = 0.05) with multiple testing corrections performed using the Benjamini-Hochberg procedure. All significant Cα peaks changed fluctations within the margin of error (< 2.5 Å). We filtered our significant Cα peaks by allowing only those observed in the non-glycosylated and glycosylated systems. This reduced the list to three potential areas in the β-barrel that could have been changed allosterically by the deletion. However, the differences in fluctuation were not consistent, meaning that the fluctuations increased when deleted in the non-glycosylated systems, while fluctuations were reduced when deleted in the glycosylated systems (Table 2). Taken together, we concluded that there is no evidence of allosteric communication within the protein caused by the deletion.

**Table 2:**
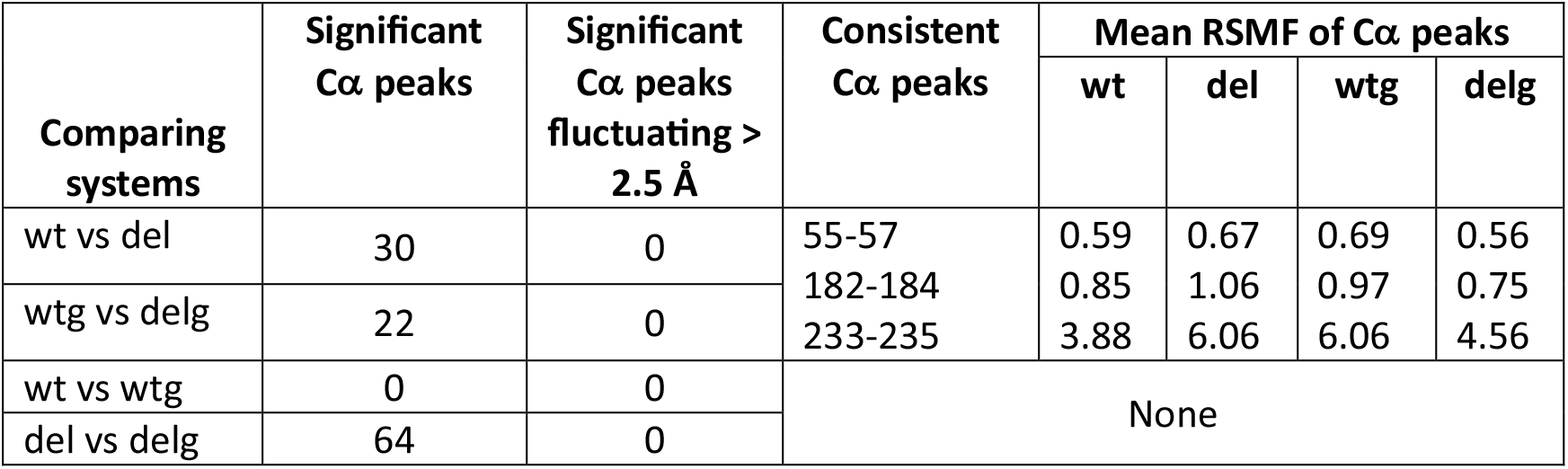
RMSF of Cα peaks (sliding window n = 3 with an increase of 1). We used the mean RMSF values from the replicates to compare different systems (column 1). Column 2 lists the number of Cα peaks that show a significant difference (p > 0.01). Column 3 lists which Cα peaks identified in Column 2 that had a difference of > 2.5 Å. Column 4 lists which Cα peaks identified in Column 2 were consistently significant across both systems, controlling for det deletion and glycosylation. Effects caused by the deletion were consistent at three positions (wt vs. del and wtg and delg comparison), listed in column 4, row 1. For the three positions, we list the mean RMSF within the replicates for each system compared in the last columns. No effects caused by the glycosylation were consistent (wt vs wtg and del vs delg), and none are listed in the table.

### The secondary structure elements are stable

We assigned secondary structure elements to the backbone for every simulation frame. We found that the 12 β-strands that comprise the β-barrel (Fig 4A) are present throughout the simulations in all four systems (Fig 4B-4C and Fig S5A-S5B). One long α-helix in the middle of the β-barrel (α-helix 2 in Fig. 4A) and an α-helix lying above the glycan (α-helix 4 in Fig. 4A) are also stable throughout the simulations. The deletion occurs within a short α-helix in a loop region (α-helix 3 in Fig. 4A). The short α-helix appears 40-60% of the time during the wt simulations (Fig 4B and Fig S5A). When the deletion is introduced, the shorter loop folds an α-helix less frequently (about 10%, Fig. 4C and Fig S5B). The deletion does not result in a loss or gain of any other β-strands or α-helices in the β-barrel. We concluded that the deletion only further destabilizes the α-helix in the deletion loop and do not cause any changes to the rest of the secondary structure elements in the domain.

**Figure 4.**
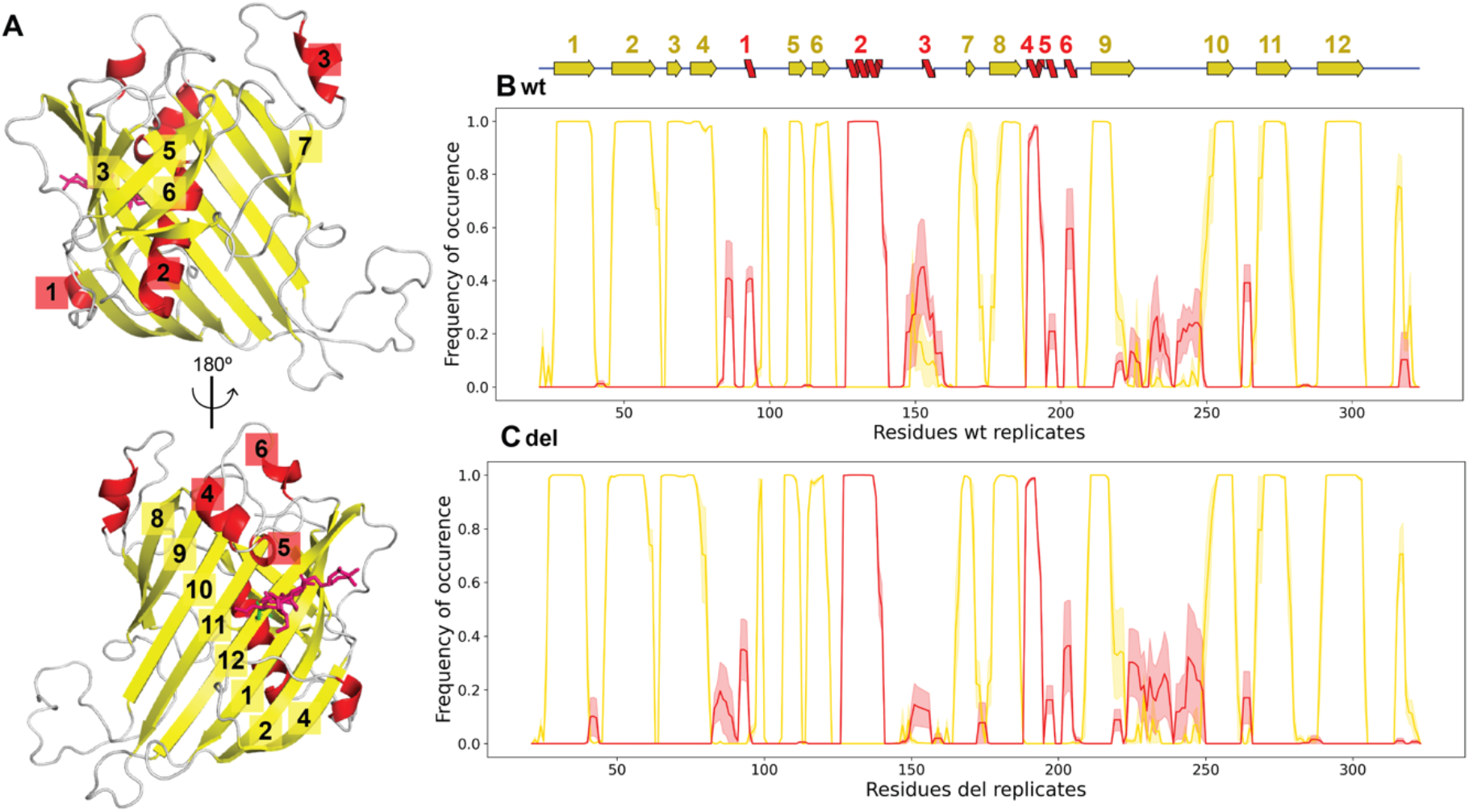
Secondary structure element assignment: **A** The 3D structure of the β-barrel is shown in two orientations (β-strands in yellow and α-helices in red). The 12 β-strands that build the β-barrel and all the α-helices are numbered. **B** The mean frequency of β-strands or α-helices (y-axis) are plotted per Cα (x-axis) of the wt system. The mean frequency is plotted as a solid line, while the ± SD values are transparent. On top of the plot is the 2D representation of the β-barrel secondary structure elements, the same as in Fig 1A, but the numbers in panel A here are included. **C** has the same plot type as panel B, but the mean frequency and SD values are from the del system.

### Principle component analysis identifies alternative conformations of a 30aa-long loop

We calculated two-dimensional principal components analysis (PCA) of the Cα atoms to identify potential differences between the systems (Fig S6A-S6C). The PCA found 19 of the 20 replicates clustered together, while a single replicate was an outlier. The outlying replicate was from the wt system and differed from the rest of the replicates throughout the simulation (Fig S6D). By extracting the frames with the highest and lowest PC1 and PC2 values from the wt replicates, we identified variable structural arrangments of a 30aa-long loop (the same flexible loop identified in the RMSF analysis). By aligning the wt replicates to the starting structure and re-calculating the RMSD for the 30aa-long loop, we found that the mean RMSD varied 7.62 Å between the wt replicates (Table S1, Fig S6E). The mean RMSD between the wt replicates for the deletion loop was far less (1.8 Å, Table S1 and Fig S6F). This showed that the separation in the PCA is caused by a lack of convergence of a highly flexible 30aa-long loop and not by the deletion loop. We repeated the PCA excluding the Cα atoms of the 30aa-long loop (aa 219-249) from all 20 replicates showing that all conditions remained clustered, and no outlier was identified (Fig 5). We concluded that most of the variance captured by the PCA did not result from the deletion.

**Figure 5.**
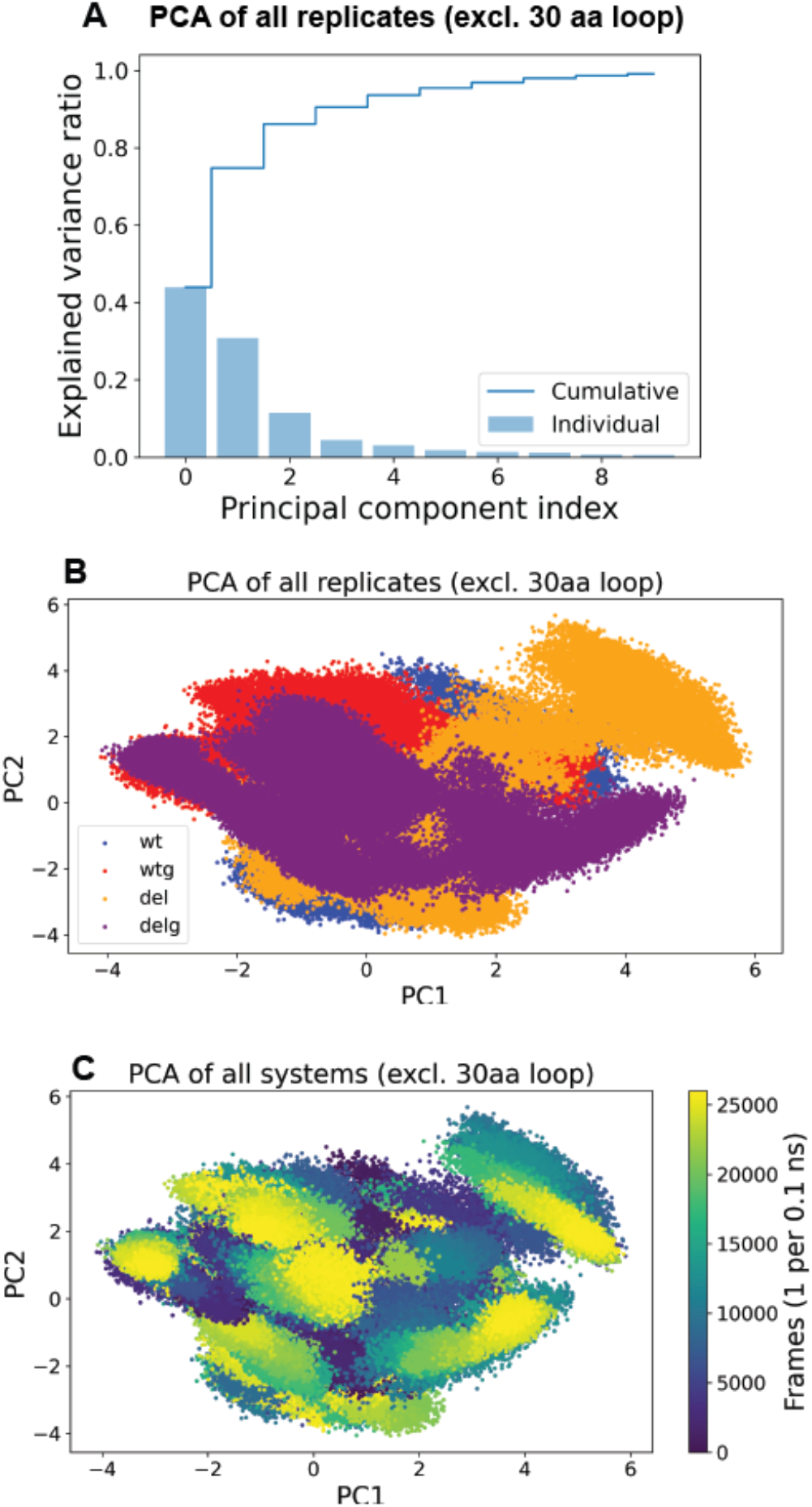
PCA: **A** The explained variance (x-axis) by PCA for 10 dimensions (y-axis). **B** PCA plot of two dimensions (x-axis: PC1, y-axis: PC2) of all 20 replicates excluding the Cα from the 30aa-long loop, wt: blue, del: orange, wtg: red, delg: purple. **C** Same plot as in panel B, but colored after frames (1 frame per 0.1 ns).

### Inhouse indel pathogenicity predictor predicts a benign deletion

To evaluate the possible pathogenicity of the deletion, we used our recently developed indel (insertions and deletions) pathogenicity predictor, IndeLLM [51]. In the absence of indel predictors applicable to variants outside of the human genome, we developed our own predictor using Protein Language Models (PLMs) [55–58]. By the use of the amino acid sequence as its only input feature, PLMs can capture complex information about protein secondary and tertiary structures across a wide range of organisms, from bacteria to humans. Benchmarking of IndeLLM showed a similar performance to other indel predictors [51]. In the context of this work, IndeLLM predicts deletion p.N153_V155del as benign. Our model was benchmarked on 7500 indels found in humans, as this organism has the most available labeled variant data. However, the pathogenicity thresholds might not transfer directly to honey bees. Therefore, we use IndeLLM to predict the pathogenicity of all possible 3aa deletions across the β-barrel (n = 300). Among all deletions, p.N153_V155del is in the top 50 (IndeLLM score: -0.48, Fig 6A-B) and above the mean IndeLLM score (-2.35), suggesting this may be a benign variant. One unique feature of IndeLLM is the option to visualize the protein structure with the impact of the indel for all amino acids in the protein since PLMs interpret the protein environment. Plotting the predicted difference between the non-deleted and deleted β-barrel per amino acids shows minor consequences for the protein, with only one amino acid in the deletion loop slightly negatively affected E147 (E147, Fig 6C-6D, -1 is damaging, while 1 is beneficial), supporting a benign variant prediction. To have a term of comparison, we visualized the lowest scoring deletion (p.E27_T29del, IndeLLM score: -13.91), which has damaging consequences across the domain, supporting a pathogenic deletion (Fig 6C and 6E).

**Figure 6.**
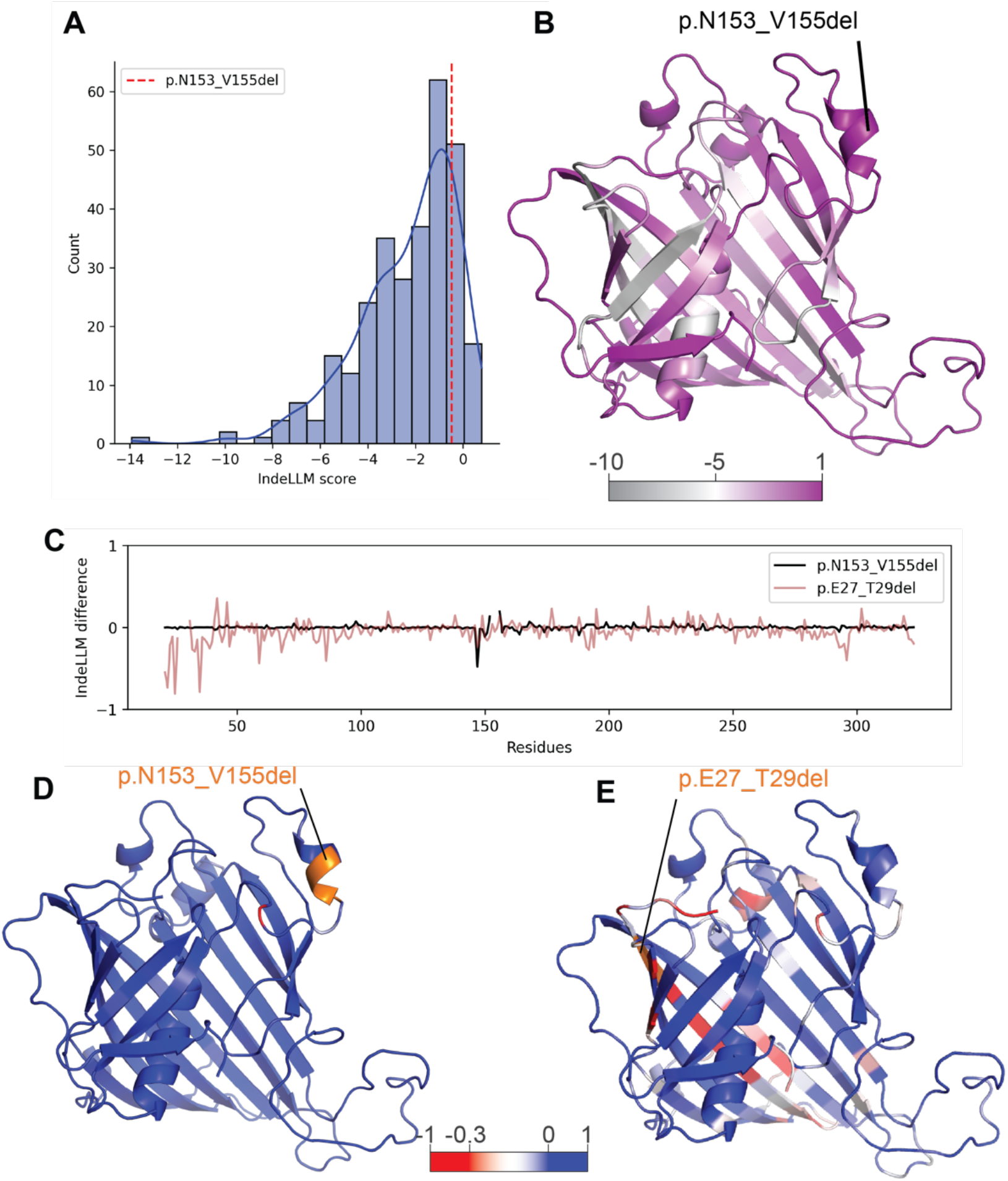
ESM: **A** The distribution of IndeLLM scores, where the deletion is colored in red. **B** The mean IndeLLM score per amino acid colored in β-barrel. The deletion is labeled. **C** The difference in IndeLLM score value (mutated value minus wt value, y-axis) for the residues in the β-barrel, for two deletions p.N153_V155 in black and p.E27_T29del in brown. **D** The difference from panel C is colored in the βbarrel structure, illustrating a low score (below -0.3) on one in the deletion loop. **E** The differences from panel C are colored for p.E27_T29del, showing several detrimental consequences for the domain.

### Glycosylation stabilizes a loop region in the β-barrel

The RMSD revealed no impact of the glycosylation on the wt or del systems. However, comparing the mean RMSF of Cα atoms in the deletion loop, we found that the glycosylation significantly reduced the flexibility of the deletion loop, but only when deleted (T-test, wt vs wtg: p = 0.706, del vs delg: p = 0.012). Although the difference in fluctuations is significant, the change in flexibility is within the margin of error (1.87 Å, Fig. S4E-S4F and Table 2). No allosteric effects were observed in β-barrel as a consequence of the glycosylation, but the glycosylation increased the occurrence of an α-helix lying above the glycan (α-helix 5 in Fig. 4A) from 20 to 40-60% (Fig S5A-S5B). The shortest distance to the α-helix is 6.3 Å (measured in the first frame of the wtg model). Taken together, the glycosylation had no major impact on the stability of the β-barrel except for introducing a more ordered arrangement in a neighboring loop region.

## Discussion

This study is the first to identify three specific deletions in the coding region of honey bee *Vg* (Fig. 1). These deletions vary widely in allele frequency, with p.S844_V845del and p.R1669del, both located outside the β-barrel, occurring at 0.06 and 0.02, respectively. In contrast, the deletion within the β-barrel, p.N153_V155del, is notably more common, exceeding 0.1 in several populations and reaching 0.5 in two. Its high frequency suggests potential biological significance. Notably, this deletion is particularly prevalent in the subspecies *A. m. mellifera*, with the highest allele frequency observed in a honey bee population on the island of Læsø, Denmark (Fig. 1B). Since 1992, a part of Læsø has, by law, been a protected mating area for *A. m. mellifera*. The genetic status or quality of the population is tested yearly using discriminant SNP markers that differentiate subspecies of European honey bees to ensure that the population is kept isolated [45,59–62]. We propose that the p.N153_V155del is specific to *A. m. mellifera* although we also detect the β-barrel deletion in the US (allele frequencies between 0.01 to 0.17). US honey bee populations is demonstrated to have a mixted genetic structure [63,64], and *A. m. mellifera* haplotypes have been detected at low frequency (0.05-0.03) [63,65]. This observation is likely due to some introduction of genetic material from European honey bees into US stocks over time [63]. The alternative explanation that the p.N153_V155del occurred independently in US honey bee stocks is theoretically possible but probably less likely.

The p.N153_V155del truncates a loop region in a highly conserved domain in Vg. We show that the deletion loop is not conserved nor close to any functional sites, indicating that the deletion loop is not directly involved in any functional roles of the β-barrel. The Rg, RMSD, and PCA analysis demonstrate that the deletion does not result in major structural rearrangements or stability loss. The consistent flexibility across all simulated conditions indicates that the deletion does not disrupt the loop’s potential for movement, and IndeLLM predicts benign consequences. The only structural consequence of p.N153_V155del is the loss of a small α-helix within the deletion loop.

Our approach for building the starting structure during the molecular dynamics simulations assumes that the β-barrel maintains its structure post cleavage. The β-barrel is documented to be proteolytically cleaved in honey bee Vg and behaves as a single functional unit [29,31]. Although our assumption might not be the case *in vivo*, we show that the β-barrel stays intact and is stable. However, the results might not be directly transferrable to the behavior of the β-barrel in the full-length honey bee Vg. In that context, the observed flexibility could be restricted by the interaction of the lipid binding site or the α-helical domain, which are structural regions close to the β-barrel (the full-length Vg is shown in Fig. 2B). However, we find that the deletion loop is positioned on the surface, at the top of the β-barrel and does not seem to interact with other regions or conserved sites in the β-barrel. Therefore, the observed flexibility of the deletion loop is highly likely to reflect the dynamics of the full-length protein as it does not seem to be restricted in any way.

The detailed molecular mechanism of the β-barrel, in protein-protein interactions or DNA binding, are poorly understood. Despite the minor structural impact we observed as a consequence of the p.N153_V155del deletion, we still acknowledge that there is a possibility that the deletion could be beneficial or alter specificity under certain conditions. It has been demonstrated that insects have unique β-barrel isoforms compared to vertebrates [31]. The insect isoform was identified in 21 insect species where the β-barrel has two additional loops. The loops were termed “insect-specific” and their preservation suggested that the loops might have a structural or catalytical role [31]. The deletion studied here, p.N153_V155del, is located in one of these insect-specific loops. In continuing research, we will attempt to link the mechanisms of the functional sites to their functional roles, such as receptor, zinc, and DNA binding. This way, we will better understand the interacting structural regions and the binding mechanisms.

The dynamic features have never been studied for any Vg chains or domain, and here, we gain new knowledge on the β-barrel’s dynamics. First, we identify a very flexible 30aa-long loop (Fig 2C). The 30aa-long loop is at the base of the β-barrel, close to the interaction site between the β-barrel and the lipid-binding cavity. The lipid-binding cavity was not included in our systems, and therefore, our system could potentially show a higher flexibility of the 30aa-long loop than what’s realistically possible in full-length Vg, considering the potential restrictions from the lipid-binding site. However, half of the 30aa-long loop is missing from the Cryo-EM model due to unresolved electron density [24], presumably caused by flexibility. This is the only region missing from the β-barrel in the Cryo-EM model, demonstrating that the 30aa-long loop is a very dynamic region also in the full-length protein. Our study shows that the 30aa-long loop is the most dynamic region in the β-barrel and therefore supports their findings. Our second interesting finding is the dynamics of the β-barrel when glycosylated. Honey bee Vg has been documented to be glycosylated on the β-barrel [24,31]. Capturing the glycan in the cryo-EM model, which is often a highly dynamic feature [66], shows that the glycan is somehow constrained. Two adjacent loops were suggested to stabilize the glycan (shown in the correct orientation of the β-barrel in Fig 2C). Here, we compare the stability of the secondary structures between the non-glycosylated and glycosylated systems and observe one loop region with an increased occurrence of an α-helix when the glycan is present (Fig 4B-4C and Fig S5A-S5B, aa 190-196). We support the claim that at least the loop region above the glycan interacts with the glycan and helps to maintain the position.

In addition to insights from molecular dynamics, we use here a novel indel predictor, which can predict consequences for the protein considering the mutation environment [51] using transfer learning. This AI-powered technique has revolutionized biological analysis, particularly protein sequence analysis, using pre-trained protein language models (PLMs) [55–58]. PLMs have shown great success in variant prediction since they are already trained on very large protein sequence datasets, capturing evolutionary and structural patterns. These models generate embeddings (numerical representations of protein sequences) that encode biochemical and structural properties and are used to capture relationships within the protein. The IndeLLM predictions shown here report the deletion as benign, both using the benchmarking threshold and performing a comparison to all possible deletions in the protein domain. The minor consequences for the protein environment of this mutation further support a neutral impact. As a negative control, we further demonstrate the ability of IndeLLM to predict pathogenicity in the domain, exemplified by p.E27_T29del, which results in damaging consequences for large regions of the domain.

We conclude that p.N153_V155del has no detrimental structural impact for honey bee Vg. Its high prevalence suggests that the deletion could be a genetic signature of a unique honey bee subspecies, *A. m. mellifera*, which increases our understanding of the endangered subspecies’ genetic pool. Our study also provides new insight into Vg’s structure-function relationships within the highly conserved β-barrel domain, highlighting its broader relevance beyond pollinators to other egg-laying animals, such as chickens and fish.

## Material and methods

### Identification of deletions

The deletions were identified using the 1086 raw consensus sequences from the *vg* sequencing dataset [38]. The sequences were aligned to the reference sequence (NCBI Gene ID: 406088) using MAFFT [67]. The deletions were identified using MEGA-X (v. 10.2.4) [68]. The allele frequencies were plotted using R (v. 2023.12.0)[69] with packages from CRAN[70] (Fig 1B-1D and Fig S1A-S1C). The p.N153_V155del was identified in 91 *A. m. mellifera* allele sequences, 1 *A. m. carnica* allele sequence, 1 *A. m. macedonica* allele sequence, and 12 *A. m*. allele sequence from the US. The haplotype sequences from *A. m. carnica, A. m. macedonica* and two from US are recombinant haplotypes [71]. The p.S844_V845del was identified in 3 *A. m*. allele sequences from the US, while p.R1669del was identified in 1 *A. m*. allele sequence from the US.

### Conservation

We identify conserved residues likely to be important for the structure and function of the β-barrel. We searched UniProt with a strict threshold likely to identify homologs of similar structure or function. The β-barrel sequence from *A. m*. (UniProt ID Q868N5, aa 21-323) was queried in Jackhmmer (v.2021_04)[72] to identify all relevant sequences from the UniProtKB. After three iterations, we selected hits with an e-value < e^-20^ (2733 sequences). Then, we applied a length filter to ensure that the selected sequences were 20% or less different in length, which reduced our sequence list to 2526. To reduce the redundancy, we used MMseq2 [73] to cluster sequences with sequence identity >=70%, resulting in 399 sequence clusters. One representative from each cluster was aligned using MAFFT, and the diversity of positions scores (DOPS)[74] was calculated to be 0.878 (with a range of 0 to 1). A DOPS score >0.80 means the MSA has high diversity, allowing robust detection of conserved residues, and the highly conserved residues may have a functional role. We considered amino acids conserved when observring a conservation socre >= 0.70. Our MSA found 15 conserved positions, shown in Fig 2A-2C.

### Preparing the starting structures

Four models of the honey bee Vg β-barrel (aa 21-323) were generated for molecular dynamic simulations: two wildtype models (wt and wtg) and two deletion models (del and delg, excluding aa 153-155). Wtg and delg include the glycan on N296. We used the Cryo-EM structure (PDB ID: 9ENS)[24] as the template structure, but the model lacks part of the 30aa-long loop region in the β-barrel (aa 232-247). Using PyMod 3.0 [75] with Modeller [76], we built a complete model identical to the cryo-EM model but included the missing residues from the 30aa-long loop region. We used the AlphaFold2 structure prediction [23] as a template for the missing loop region, and Modeller to get a refine the model. The resulting model is a complete structural representation of the β-barrel, the wt model. The glycan on N296 was imported from the cryo-EM model to build the wtg model. The del model was built using the wt model. We used ColabFold [77] to generate a β-barrel excluding aa 153-155 to have a structural prediction of a shorter loop. Next, we removed 153-155 from the wt model sequence. Using Modeller (as described above), we generated a homology model using the wt model and the AlphaFold2 predicted deletion loop region (aa 150-157) as a template. The resulting model is a structural representation of the β-barrel with the documented deletion, the del model. The glycan on N296 was exported from the cryo-EM model to build the second deletion model (delg). We renumbered the aa in the del models to 21-152 and 156-323 so that the aa numbering in the wt, wtg, del, and delg models are identical, and the wt and wtg are the only models including aa 153-155.

### System preparation and molecular dynamics simulations

The systems were prepared using CHARMM-GUI [78–80]. We used the Solution Builder for the non-glycosylated systems (wt and del) and Glycan Reader & Modeler [81–83] for the glycosylated systems (wtg and delg). The minimization, equilibration, and production run were done using Amber 22 (see the method section in the supplementary data for more information). The input and output files are provided in the supplementary information.

### Trajectory analysis

We simulated 3 μs per replicate, saving every 100 ps, resulting in 30,000 trajectory frames. After production, we combined the trajectory files and removed the water molecules. The cleaned trajectory files were used for the following calculations. For each replicate, the RMSD, RMSF, and Rg were calculated on the Cα atoms using MDTraj [84]. The secondary structure assignment was also done with MDtraj using all protein atoms. We combined the Cα from the cleaned trajectories (excluding aa 153-155 from wt and wtg systems to ensure equal number of atoms) to calculate PCA using Scikit-learn [85]. We extracted the top two principal components (PC) that explained most of the variance of the data (Cumulative explained variance = 0.34 %; see Fig S6C for the explained variance of 10 components). We extracted PDB files of the specific frames in which the PC values were more extreme. The frames z were 8674 (Max PC1 6.30) and 7118 (Min PC2 -0.42) for wt1, 24888 (Min PC1 -6.05) and 1931 (Min PC2 -2.21) for wt2, 3759 (Min PC1 -6.43) and 202 (Min PC2 -1.60) for wt3, 921 (Max PC1 7.44) and 3330 (Min PC2 -1.46) for wt 4 and 1914 (Max PC2 8.05) for wt5. The frames were aligned and analyzed in PyMol [86]. In the second PCA (excluding the 30aa-long loop), the cumulative explained variance of the first two components was 0.75% (see Fig 5A for the explained variance of 10 components). The supplementary information provides all the processing and plotting scripts.

### IndeLLM prediction

We used the IndeLLM source code [51] to predict the impact of all possible 3 aa deletions on the β-barrel domain of Vg. The deletions were generated by using a sliding window of size 3 with a step of 1, generating a total of 300 sequences with deletions, and obtaining an IndeLLM score per sequence.

## Acknowledgements

We greatly thank Maria Laura De Sciscio (University of Rome) for advices on on molecular dynamics protocols, and Carlos Henrique Bezerra Da Cruz (University College London) for reading and providing comments on the manuscript. We also thank Ole Kilpinen and Flemming Vejsnæs (Danish Beekeeping Association) for reading and commenting on the manuscript. The authors acknowledge The Research Council of Norway grant numbers 335244 and 350231 for funding toward running costs, travel grants, and conference support.

## Author contributions

Conceptualization, VL, GVA; methodology, OGIC, FF, CO; validation, OGIC, FF, CO, GVA; formal analysis, VL, OGIC; investigation, VL; resources, FF, GVA; data curation, VL; writing—original draft preparation, VL; writing—review and editing, OGIC, FF, CO, GVA; visualization, VL; supervision, FF, GVA; project administration, VL; funding acquisition, FF, GVA All authors have read and agreed to the published version of the manuscript. All authors reviewed the manuscript.

## Supplemental information

Figure S1 – homozygotic honey bees for the identified deletions. Figure S2-S3 and Table S1 replicate RMSD analysis, Figure S4 replicate RMSF analysis, Figure S5 DSSP analysis for glycosylated system and Figure S6 detailed PCA analysis. Method description for molecular dynamics simulations.

## Resource availability

The starting models, input and output files from the system preparation, minimization, equilibration, and production, as well as the Python scripts generated and analyzed during the molecular dynamics in our study, are available in the supplementary information MD_Analysis.zip and last section on the supplementary data.

The authors declare no conflicts of interest.

