## Supplementary Material for "Assessing structure-function impacts on Vitellogenin by leveraging allelic variant occurring in honey bee subspecies *Apis mellifera meliffera*"

### Supplementary data

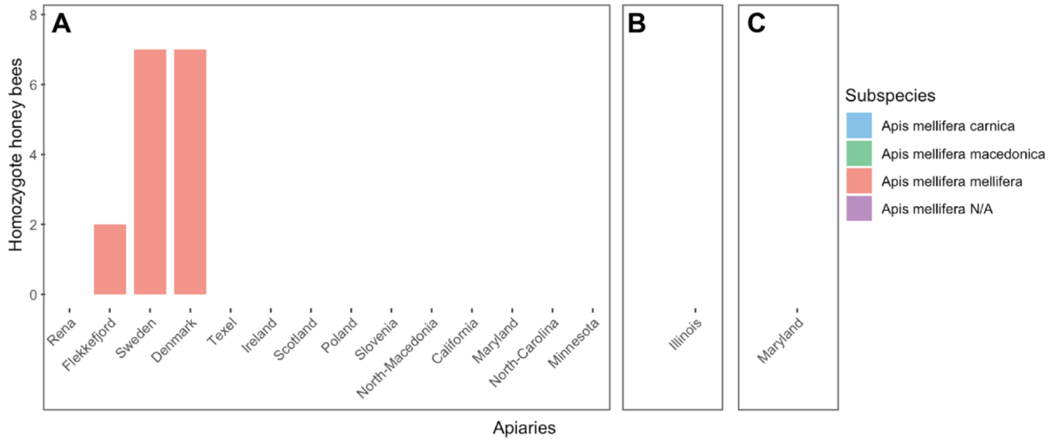


***Figure S1 Homozygotic honey bees: A** The identified homozygotic bees for p.N153_V155del. The number of honey bees (y-axis) is plotted per apiary (x-axis). The apiaries are colored by subspecies (legend). **B-C** No homozygotic bees were identified for p.S844_V845del (B) or p.R1669 (C).*

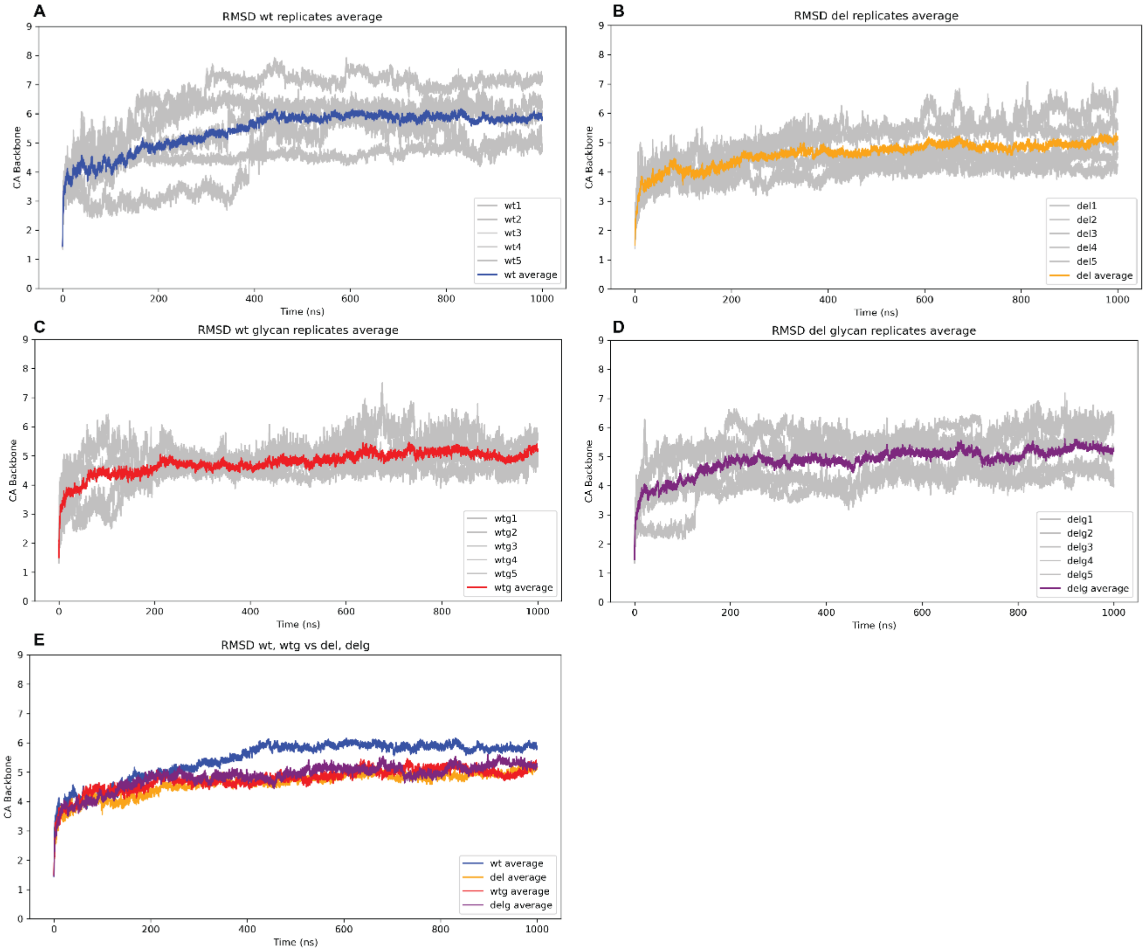


***Figure S2 RMSD after 1µs** The RMSD between the Cα atoms (y-axis) over time (x-axis) for each replicate are plotted in grey, while the mean RMSD is plotted in colors. **A** wt replicates and mean (blue). **B** del replicates and mean (orange). **C** wtg replicates and mean (red). **D** delg replicates and mean (purple). **E** Plott of all mean RMSD values from all systems.*

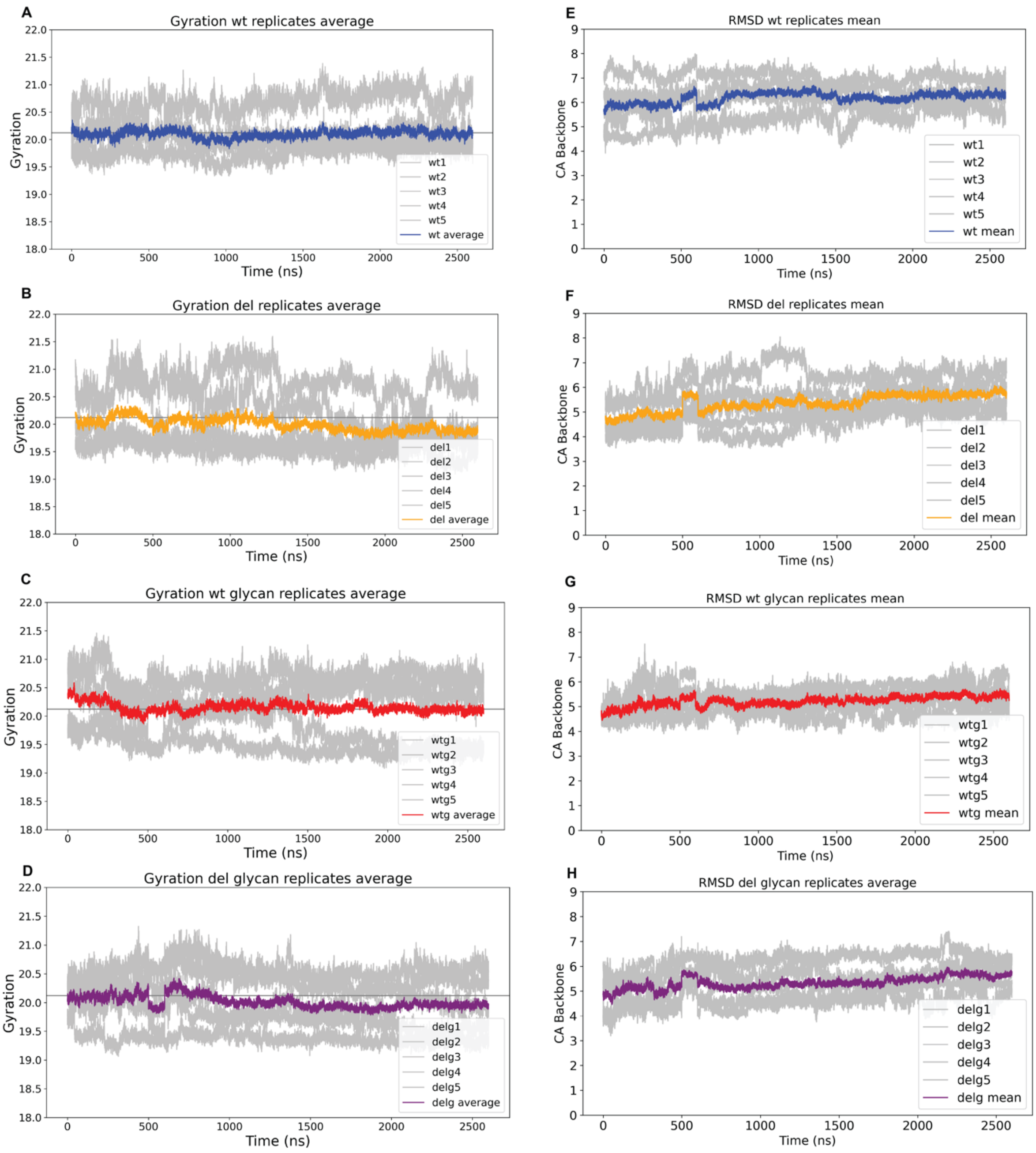


***Figure S3 Rg and RMSD after 3μs for all replicates: A-D:*** *The Rg (y-axis) over time (x-axis) for each replicate are plotted in grey, while the mean Rg is plotted in colors. The grey horizontal line shows the Rg of the starting wt model (20.12 Å).* ***E-H*** *The RMSD for the Cα atoms (y-axis) over time (x-axis) for each replicate are plotted in grey, while the mean RMSD is plotted in colors.* ***A and E*** *wt replicates and mean (blue).* ***B and F*** *del replicates and mean (orange).* ***C and G*** *wtg replicates and mean (red).* ***D and H*** *delg replicates and mean (purple).*

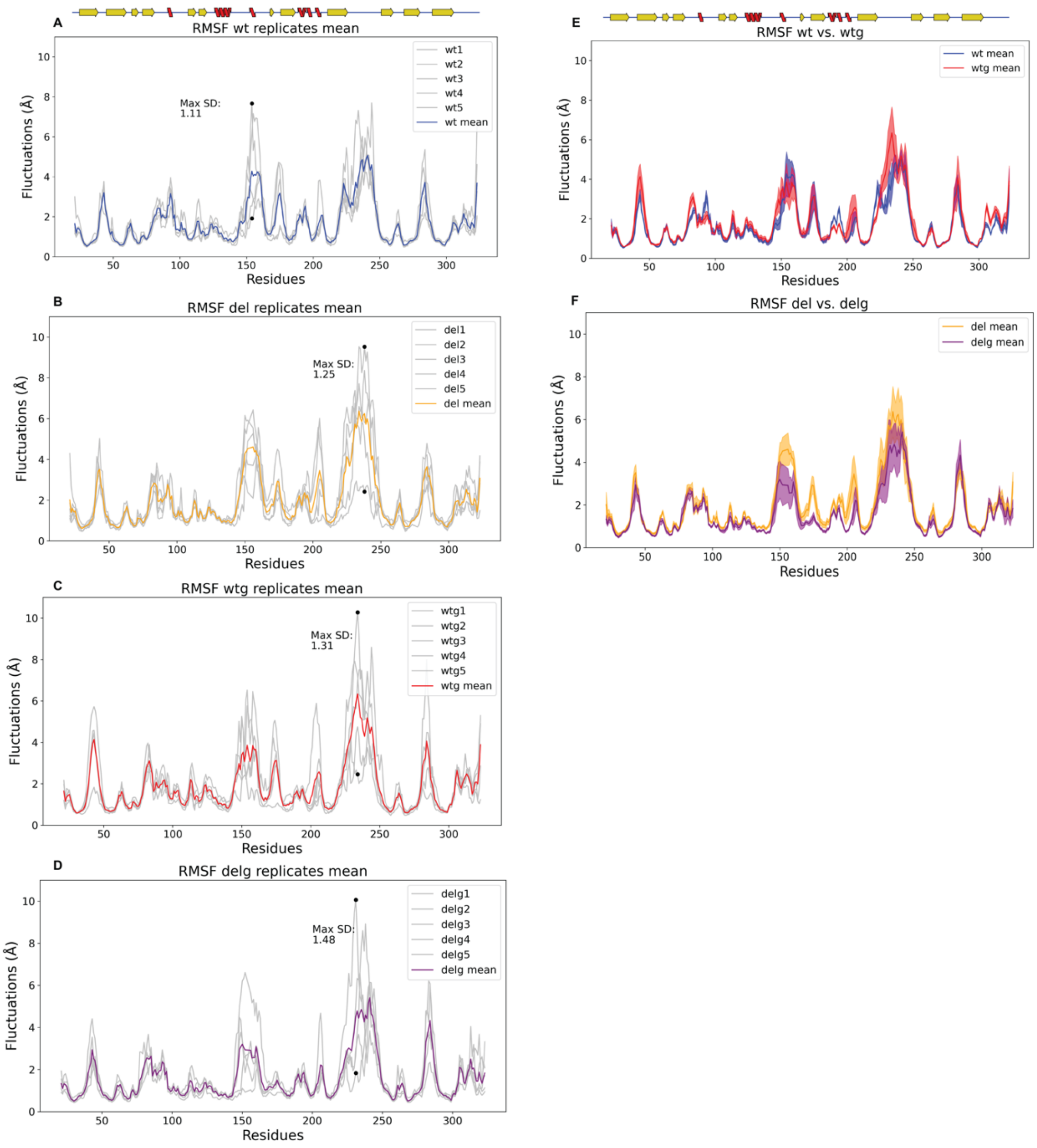


***Figure S4 RMSF after 3µs for all replicates and analysis: A-D:*** *The RMSF (y-axis) for the Cα atoms (x-axis) for each replicate are plotted as separate grey lines, while the mean RMSF is plotted in colors. The maximum SD observed between the replicates is plotted as black dots.* ***A*** *wt replicates and mean (blue).* ***B*** *del replicates and mean (orange).* ***C*** *wtg replicates and mean (red).* ***D*** *delg replicates and mean (purple).* ***E*** *The mean RMSF (y-axis) for each Cα in the wt (blue) and wtg (red) systems (x-axis, aa 21-323). The mean values are plotted as a solid line, while the ± SD values are transparent. On top of the plot is the 2D representation of the β-barrel secondary structure elements, the same as in Fig 1A.* ***G*** *has the same plot as in panel F, but the mean RMSF values are from the del (orange) and delg (purple) systems.*

1

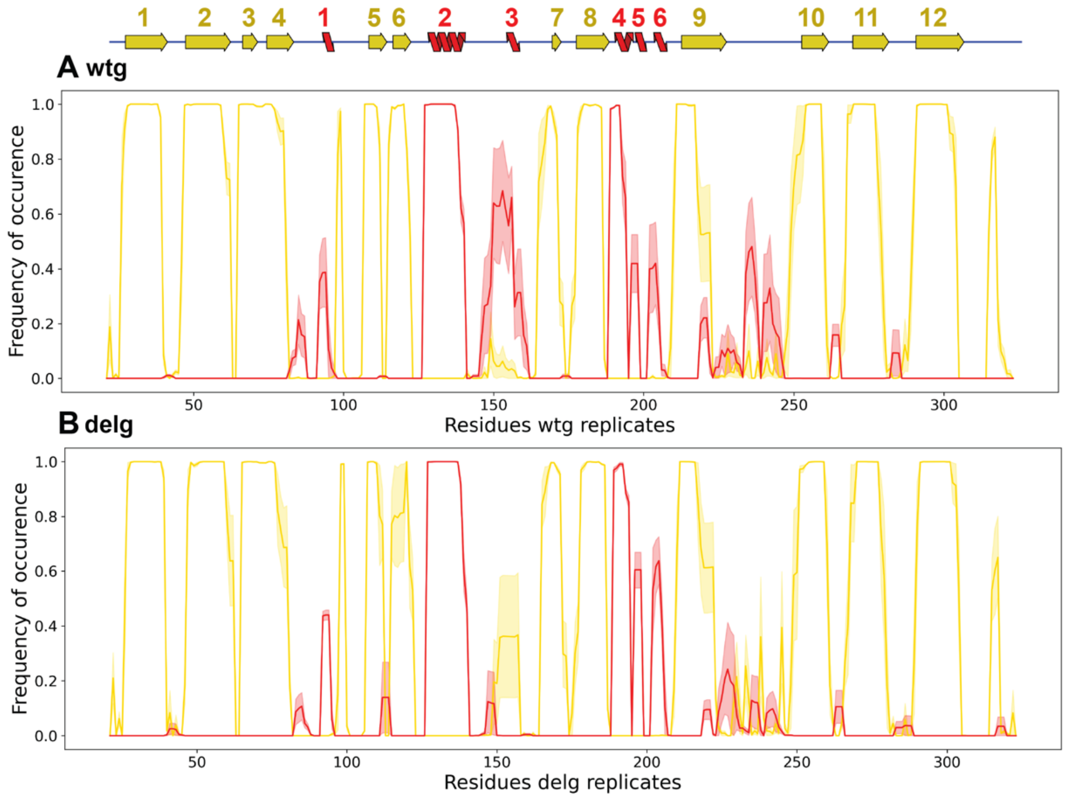


***Figure S5 Secondary structure element assignment for the glycosylated systems A*** *The mean frequency of β-strands or α-helices (y-axis) are plotted per Cα (x-axis) of the wtg system. The mean frequency is plotted as a solid line, while the ± SD values are transparent. On top of the plot is the 2D representation of the β-barrel secondary structure elements (β-sheets in yellow and α-helices in red).* ***B*** *is the same plot as in panel A, but the mean frequency and SD values are from the delg system.*

2

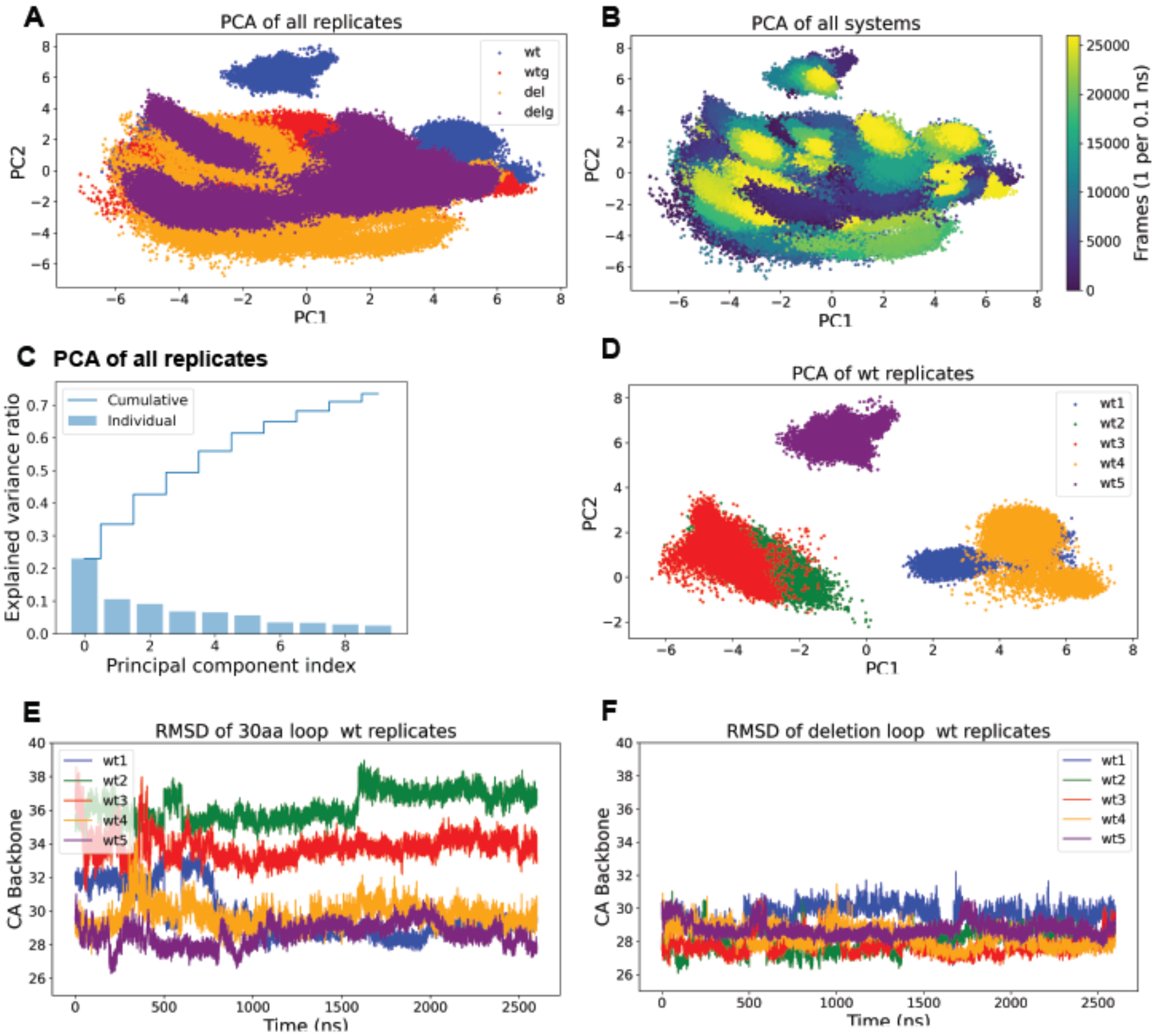


***Figure S6 PCA: A** PCA plot of two dimensions (x-axis: PC1, y-axis: PC2) of all 20 replicates, wt: blue, del: orange, wtg: red, delg: purple. **B** Same plot as in panel A, but colored after frames (1 frame per 0.1 ns). **C** The explained variance (x-axis) by PCA for 10 dimensions (y-axis). **D** Same plot as in panel A, but only includes the wt replicates: 1 (blue), 2 (green), 3 (red), 4 (orange), and 5 (purple). **E** The RMSD for the Cα atoms of the 30aa-long loop when (y-axis) over time (x-axis) for each wt replicate with the same colors as in panel D. **F** Same plot as in panel E, but the mean RMSD for the deletion loop.*

3

| Replicates | | Deletion loop (aa 141-167) | 30aa-long loop (aa 219-249) |
|---|---|---|---|
| wt1 | Overall mean | 29.54 | 29.73 |
| | Range (Å) | 26.55-32.21 | 27.58-33.86 |
| | SD | 0.72 | 1.45 |
| wt2 | Overall mean | 28.03 | 36.23 |
| | Range (Å) | 26.09-31.03 | 33.85-38.96 |
| | SD | 0.64 | 0.85 |
| wt3 | Overall mean | 27.74 | 33.82 |
| | Range (Å) | 26.51-30.61 | 31.17-38.58 |
| | SD | 0.56 | 0.75 |
| wt4 | Overall mean | 28.25 | 29.74 |
| | Range (Å) | 26.81-31.48 | 27.44-34.62 |
| | SD | 0.60 | 0.72 |
| wt5 | Overall mean | 28.74 | 28.61 |
| | Range (Å) | 27.60-30.79 | 26.28-31.02 |
| | SD | 0.45 | 0.68 |
| wt1-5 | Overall mean | 28.46 | 31.62 |
| | Range (Å) | 27.42-29.56 | 30.52-33.46 |
| | SD | 0.25 | 0.34 |

***Table S1:*** *RMSD calculations for the deletion loop and the 30aa-long loop after superimposing the full-length fold. Rows 1-5 show the calculations for RMSD for each wt replicate. For the two loop regions (columns 3 and 4), we calculate the overall mean for all C$\alpha$, the range of difference in Å, and the standard error of the mean (SD). The last row includes the same calculations for RMSD, but we are comparing all the wt replicates using the mean values.*

**Methods: Molecular dynamics simulations**

The PDB files for each system were uploaded to generate a PDB in CHARMM-GUI [1–3]. The waterbox size was selected by default (Rectangular with an edge distance of 10.0 Å). Ions were added using the default options (Monte-Carlo, KCl) to solvate the molecule. The grid information was generated automatically using the Particle Mesh Ewald Fast Fourier Transform algorithm (PME-FFT). We used the AMBER FF19SB force field [4] for proteins, the GLYCAM_06j force field [5] for the glycan and TIP3P [6,7] for water.

Minimization was run for all systems with 5000 maximum cycles; the first 2500 cycles used the steepest descent algorithm. Positional restrains (1 kcal $mol^{-1}$ $angstrom^{-2}$) were placed on all protein and glycan atoms (wt: 1-303, wtg: 1-308, del: 1-300, and delg: 1-305). Following minimization, the systems were equilibrated under the NVT ensemble, raising the temperature stepwise to 300K (50K, 100K, 200K, and 300K) with 125 ps per step, using the same positional restraints as during the minimization.

The production simulations were run under the NPT ensemble, with temperature control at 300K using Langevin dynamics (friction coefficient at $1ps^{-1}$), potential energy control (nonbonded cutoff at 9 Å), and pressure control using Montecarlo Barostat (isotropic with target pressure at 1 bar). The SHAKE [8] algorithm constrained all protein hydrogen bonds, while water molecules were treated with SETTLE [9]. The systems were simulated stepwise for 3µs (100 ns per step). The energies, coordinates, and restart files were printed every 100 ps.

The minimization step was performed with the primary molecular dynamics engine pmemd implementation in AMBER 22 [10], while the equilibration and production runs were performed with single precision pmemd.cuda_SPFP in AMBER 22.
